# HIV superinfection reveals sequential reservoir reactivation and immune-driven rebound dynamics

**DOI:** 10.64898/2026.03.18.712773

**Authors:** F. Harrison Omondi, Aniqa Shahid, Natalie N. Kinloch, Winnie Dong, Maggie C. Duncan, Fatima Yaseen, Evan Barad, Nadia Moran-Garcia, Amanda Cabral da Silva, Vitaliy Mysak, Don Kirkby, Mario Ostrowski, Rebecca M. Lynch, Chanson J. Brumme, Colin Kovacs, R. Brad Jones, Guinevere Q. Lee, Zabrina L. Brumme

## Abstract

Superinfection, where a person with HIV acquires a second phylogenetically distinct strain, provides an opportunity to study reservoir evolutionary dynamics because the two strains can be tracked over time. We here characterize such a case that originally went undetected by clinical monitoring. The participant, who expressed the protective HLA-B*57:03 allele, initially controlled his subtype B infection but lost control after superinfection with a unique recombinant form (URF), which came to dominate in plasma without displacing the original B strain. Five years after initiating therapy, replication-competent proviruses from both strains persisted in blood, though URF proviruses dominated (80%). Despite their unequal reservoir representation, both strains rapidly rebounded upon treatment interruption, though most rebounding sequences belonged to a URF subclade that predated ART initiation. During the treatment interruption, genetically distinct viral lineages continually appeared in plasma, consistent with sequential “waves” of reactivation of diverse reservoir clones. Notably, immune escape variants emerged from the reservoir in a later “wave”, displacing initial susceptible variants and underscoring the important role of immune responses in shaping rebound dynamics. Sequence analysis revealed that the URF transmitted/founder virus already harbored key HLA-B*57:03-associated escape mutations, likely explaining the loss of viral control after superinfection. Our results reveal that superinfection can remain undetected despite routine HIV clinical monitoring, that pre-adaptation of the superinfecting strain to host HLA can undermine immune control, that rebound virus can originate from reservoir clones predating ART initiation, and that immune responses can actively shape rebound dynamics by driving escape variant outgrowth during treatment interruption.

**IMPORTANCE:** This case demonstrates that HIV superinfection can go undetected by routine clinical monitoring, underscoring the importance of assays that capture HIV diversity — particularly those used to screen participants for cure trials. It also illustrates how pre-adaptation of a superinfecting strain to host HLA-restricted immune responses can evade existing immunity, with implications for understanding natural immune control, and underscoring the challenge of viral diversity for vaccine and immunotherapy design. The sequential emergence of genetically diverse HIV lineages during treatment interruption, including minority immune escape variants that emerged from the reservoir to displace the initial susceptible population, highlights the difficulty of predicting rebound virus composition and underscores the key role of immune responses in shaping rebound dynamics. Critically, escaped rebound variants can then reseed the reservoir, thereby enriching it in escaped forms, highlighting a specific vulnerability for cure approaches that harness natural immune responses to eliminate reservoir cells.

## INTRODUCTION

Human Immunodeficiency Virus (HIV) infection cannot currently be cured because intact, replication-competent copies of the virus genome persist in long-lived memory CD4+ T cells as integrated proviruses (1–4). These cells, called HIV reservoirs, can reactivate at any time to produce infectious virus (5, 6), and for this reason antiretroviral therapy (ART) must be maintained for life. Much of our understanding of when individual within-host HIV sequences become archived in the reservoir, how long they persist there, and which are the first to re-emerge when ART is interrupted, comes from within-host HIV evolutionary studies that identify the relative age of each provirus based on when it, or sequences closely related to it, circulated in the host (7–11). Cases of superinfection, where a person with HIV acquires a second, phylogenetically distinct infection, provide a similar opportunity to study reservoir establishment and rebound dynamics, as the two distinct strains can be molecularly tracked over time (12).

Though superinfection is estimated to occur at rates up to 8% depending on the population (13–15), cases remain inherently challenging to identify as no routine tests reliably detect them (16). Longitudinal HIV drug resistance genotyping combined with phylogenetic monitoring represents one such opportunity, though this approach best identifies cases where the superinfecting strain outcompetes the original one, prompting a clear re-positioning of the individual’s sequences in the phylogeny. Cases where both original and superinfecting strains continue to co-exist can be more challenging to detect, particularly when performing bulk sequencing, which yields a composite sequence featuring nucleotide mixtures at positions exhibiting within-host diversity. Variants present at frequencies below 20% may be missed, as bulk sequencing cannot reliably detect variants below this threshold (17), while samples with large numbers of nucleotide mixtures may fail quality controls and not be reported. While next-generation sequencing (NGS) better detects within-host variation, all sequencing approaches nevertheless require the upstream amplification method to capture both strains, for individuals to be followed longitudinally, and for metrics to be in place to distinguish superinfection from normal within-host viral evolution (16, 18–20). As a result, superinfection is often identified serendipitously.

We retrospectively characterized a case of HIV superinfection that went undetected by routine clinical monitoring, that was discovered after the participant enrolled in a research study to characterize their HIV reservoir. Analysis of *env* genetic diversity in viruses cultured from CD4+ T cells during ART revealed two distinct HIV strains, one subtype B and the other distinctly non-B (21), an observation that was confirmed by two independent laboratories (22). Here, we analyzed HIV diversity in additional samples, including plasma collected in early infection and again just prior to ART initiation, blood cells collected during ART, and longitudinal plasma collected following ART interruption. Leveraging each strain’s distinctive genetics, we determined the order in which each strain established infection, examined the composition of the proviral landscape during ART, and characterized the evolutionary dynamics of rebound viremia.

## RESULTS

### Participant clinical and sampling history

The participant, a male in his 50s of African descent, was diagnosed with HIV in mid-2010, after likely acquiring the virus during a self-reported high-risk encounter earlier that spring (an HIV test performed in early 2010 was negative) (**Figure 1**). His baseline viral load, recorded in Fall 2010, was 3.1 log_10_ HIV RNA copies/mL. This relatively low viral load is consistent with initial viremic control, likely attributable at least in part to his carriage of the protective HLA-B*57:03 allele (23, 24). By mid-2012 however, his plasma viral load had reached 4.9 log_10_ HIV RNA copies/mL, and he initiated ART containing emtricitabine, tenofovir and raltegravir. A clinical drug resistance genotype performed at this time confirmed a lack of drug resistance and surprisingly classified the HIV subtype as B, even though superinfection had already occurred (see below). Of note, clinical coreceptor usage genotyping performed at this time reported CCR5-using HIV with an envelope V3 loop sequence of C[A/V]RPNNNTRKSIRIGPGQTFYATNDIIG[N/D]IRQAHC. Though coreceptor genotyping is not used to determine HIV subtype (due to the short length and limited phylogenetic signal of the V3 loop), this sequence is distinctly non-B. Aside from an isolated viral blip in 2013 to 68 copies/mL, the participant maintained viral suppression until mid-2018 when he briefly interrupted ART. Following this, HIV re-emerged in plasma within two weeks. The ART interruption lasted less than two months, and suppression was re-achieved within two weeks.

**Figure 1.**
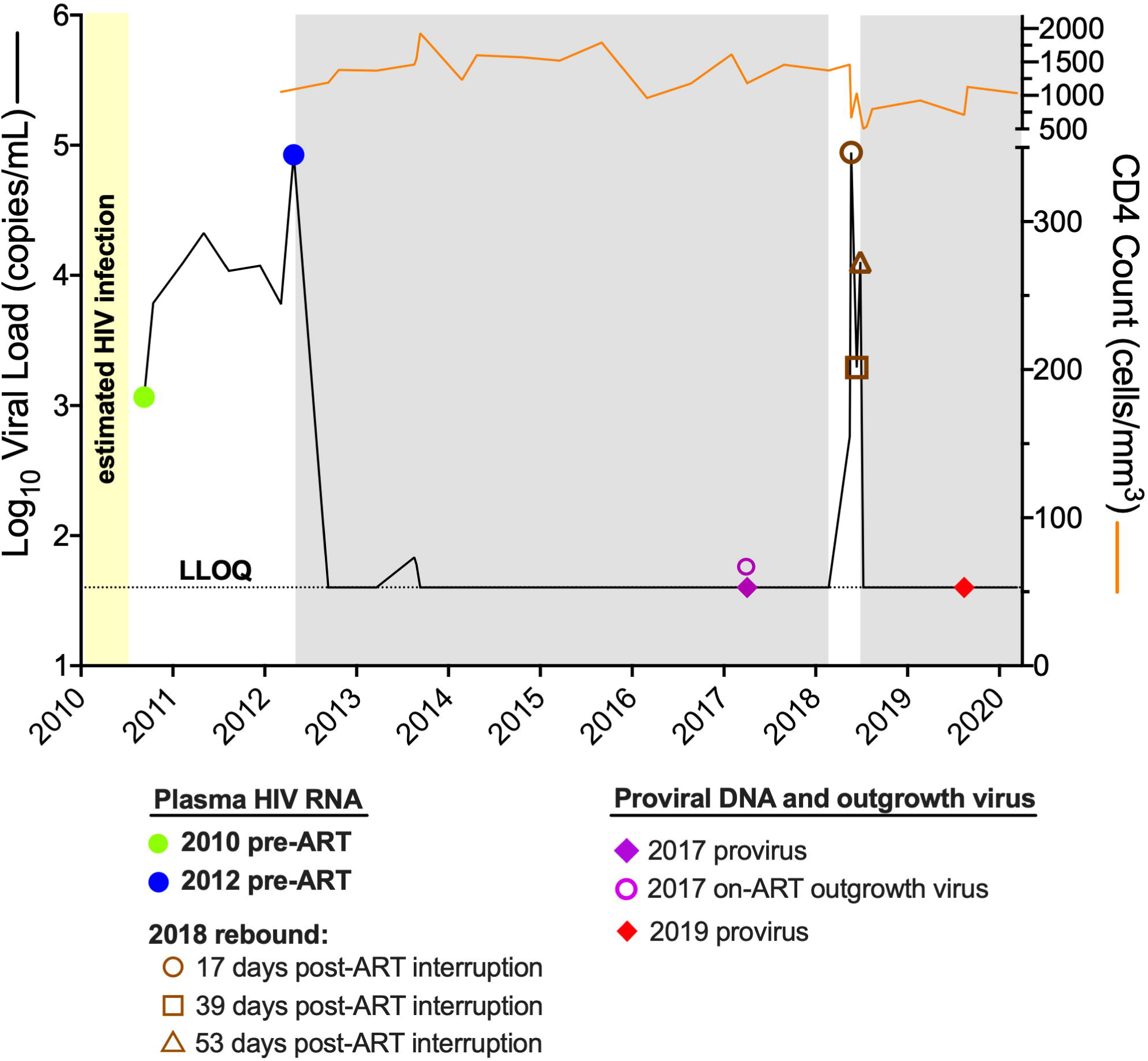
Participant clinical history and sampling timeline. CD4 counts (orange solid line) and plasma viral loads (solid black line) are shown longitudinally, where the dotted horizontal represents the viral load assay lower limit of quantification (LLOQ). Grey shading denotes ART. Symbols show pre-ART plasma HIV RNA sampling (green and blue circles), on-ART blood proviral DNA sampling (purple and red diamonds), on-ART viral outgrowth sampling (open purple circle) and plasma HIV RNA sampling during the treatment interruption (open brown symbols).

### Retrospective confirmation of superinfection

Retrospective single-genome amplification of *gp120* (25 sequences) and *pol* (31 sequences) from baseline plasma collected in 2010 revealed exclusively subtype B HIV (**Figures 2 and S1**), indicating the initial infection was subtype B. Overall genetic diversity was limited, consistent with infection only three months prior. By contrast, single-genome amplification of *gp120* (76 sequences) from plasma collected in 2012 revealed descendants of the original subtype B virus along with sequences belonging to a genetically distinct unique recombinant form (URF) (**Figure 2**). By this time, the URF dominated in plasma (90% of sequences) and exceeded the subtype B sequences in terms of within-host genetic diversity, even though it was acquired later (the median patristic distance separating URF sequences was 0.0127 compared to 0.0116 for the B strains). The URF was therefore the superinfecting strain, which became dominant without displacing the original subtype B population. Though it is not possible to estimate the superinfection date from sequences sampled at a single timepoint, the diversity of the URF sequences suggested that superinfection was not particularly recent. There was no evidence of within-host recombination between B and URF sequences in the tree (**Figure 2 inset**), nor when analysing within-host sequences for recombination using RDP4 (25) (not shown). Superinfection with the URF, and lack of within-host recombination, was further corroborated by analysis of 34 single-genome *pol* sequences from the 2012 plasma sample (**Figure S1**).

**Figure 2.**
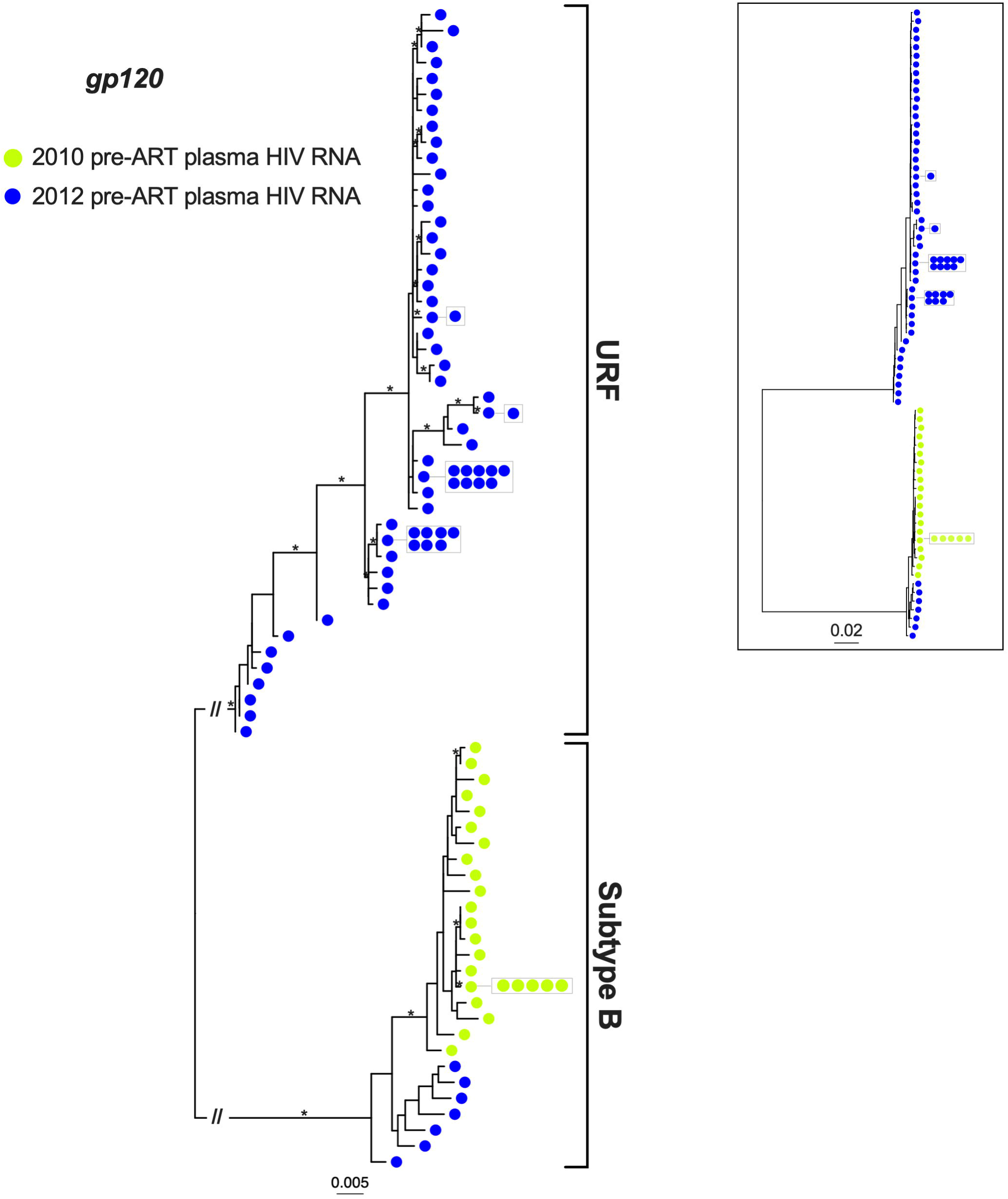
Maximum likelihood within-host phylogeny of *gp120* sequences sampled pre-ART. Colored circles denote distinct pre-ART plasma HIV RNA sequences sampled in 2010 (green) and 2012 (blue), with identical sequences shown in adjacent boxes. Asterisks identify branches with bootstrap support of ≥90%. Scale is in estimated substitutions per nucleotide site. The tree is midpoint-rooted, with the long subtype-specific branches truncated with a “//” symbol to save space. The inset shows the full tree with untruncated main branches.

### Both initial and superinfecting strains were preserved in the reservoir, with the latter dominating

From a sample collected in 2017, after ∼5 years of suppressive ART, we isolated 196 near-full-length proviruses (20% subtype B, 80% URF) from blood CD4+ T cells by single-genome sequencing, and an additional two (one each from subtype B and the URF) by limiting-dilution viral outgrowth (**Figure 3A**). This confirmed that both strains were preserved in the reservoir, at comparable frequencies to their respective plasma virus populations just prior to ART (**Figure 3A**). As is typical during long-term ART (26, 27), proviruses from both strains were predominantly defective, featuring hypermutation, 5’ leader defects, or large deletions. Nevertheless, 26 (13%) of all proviruses, a total that included 22 URF and two subtype B proviruses recovered from CD4+ T cells, plus the two viral outgrowth isolates (one each per subtype), were genetically intact. This allowed us to confirm the URF as a complex mosaic of subtype G and CRF02_AG that has not been previously described (**Figure S2**). We also observed proviruses that were genetically identical over the sequenced region (**Figure S3**), consistent with clonal expansion (28–32). In total, 63 proviruses (57 URF, 6 subtype B sequences represented by hatched pattern), representing 32% of all proviruses sampled, were members of a clonal set. The three largest clones, all belonging to the URF, were an intact provirus (observed 15 times), a provirus with its 3’ half deleted (observed 12 times), and a hypermutated provirus (observed 9 times) (**Figure S3)**. Consistent with pre-ART clinical genotyping, all proviruses recovered during ART were CCR5-using, and none contained drug resistance mutations. Also consistent with analysis of subgenomic plasma HIV sequences collected prior to ART, no proviruses represented B/URF recombinants.

**Figure 3.**
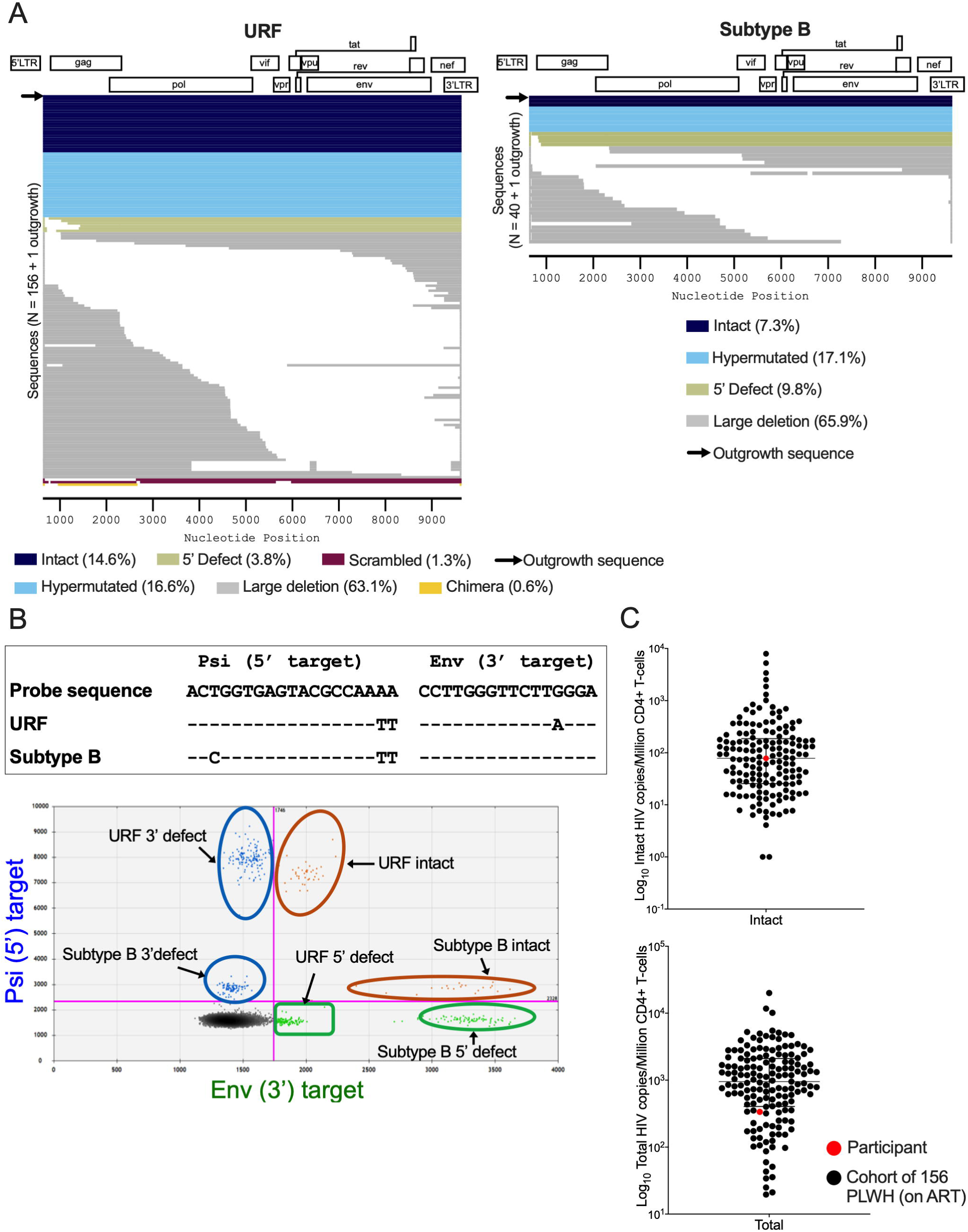
Proviral composition during ART. (A) Near-full-length URF (left panel) and subtype B (right panel) proviral genomes isolated during ART, colored by genomic integrity, with the frequencies of each category shown below each panel. “Scrambled” denotes proviruses with one or more HIV gene regions that are out of order; “Chimera” denotes proviruses with embedded human genome fragments (B) Top: Alignment of the URF and subtype B consensus sequences in the IPDA probe region, where hyphens (-) indicate matches. Bottom: IPDA plot showing the impact of subtype-specific polymorphisms on signal amplitude. (C) Top: Intact HIV copies per million CD4+ T cells in the participant (red circle) and 156 individuals receiving ART who had not experienced superinfection (black circles). Bottom: Same plot, but for total HIV copies per million CD4+ T cells. Line and whiskers show median and interquartile range.

The participant’s two HIV strains produced different signal amplitudes in the intact proviral DNA assay (IPDA) due to polymorphism in the probe-binding regions, allowing us to estimate overall and strain-specific reservoir sizes (**Figure 3B**). Here, one challenge was the URF’s G-to-A polymorphism at position 13 of the *env* probe, which dramatically reduces signal amplitude and makes it difficult to distinguish small numbers of *env*-positive events from the large double-negative population when typical (*e.g.* ∼250,000) cell numbers are assayed (22). To overcome this, we assayed DNA from 1.7 million CD4+ T cells. The IPDA revealed an overall intact proviral burden of 78 copies/million CD4+ T cells, and an overall total proviral burden of 335 copies/million CD4+ T cells (**Figure 3C**). For context, these measurements represented the 50^th^ and 21^st^ percentiles, respectively, of a cohort of 156 individuals on ART who had not experienced superinfection, indicating that HIV superinfection does not necessarily produce a higher-than-average reservoir or total proviral loads. Consistent with the sequencing results, the URF dominated: intact and total proviral URF burdens were 50 and 201 copies/million CD4+ T cells, respectively, while corresponding measurements for subtype B were 28 and 134 HIV copies/million CD4+ T cells, respectively (**Figure 3B)**. Of note, the IPDA-estimated intact proviral loads substantially exceeded the QVOA-estimated reservoir size, which was 0.27 infectious units per million CD4+ T cells. Since the IPDA estimates genomic intactness based on only two HIV targets, it can overestimate reservoir size if many proviruses have defects outside the target regions (33, 34), but our provirus’ sequencing showed that this was not the case presently. It is therefore more likely that the QVOA only activated a minority of genetically-intact proviruses (35). Regardless, the *in vitro* outgrowth of both B and URF confirmed that replication-competent representatives of both strains were preserved in the reservoir.

Phylogenetic analysis of on-ART proviral *gp120* diversity in context of pre-therapy plasma HIV RNA diversity yielded additional insights (**Figure 4**). One of the intact subtype B proviruses as well as the subtype B outgrowth virus branched in-between the 2010 and 2012 subtype B plasma viruses with strong branch support (**Figure 4** near bottom of tree), suggesting they arose sometime between those dates. Two subtype B proviruses clustered with 2010 plasma viruses, demonstrating that proviral seeding into long-lived infected cells begins immediately following transmission (36–38), though the fact that both of these proviruses were defective suggested that few *replication-competent* proviruses dating to transmission persisted after five years on ART. Three intact URF proviruses clustered among the plasma sequences sampled in 2012 (**Figure 4**), suggesting that these arose immediately pre-ART. All other intact URF proviruses, as well as the URF outgrowth sequence, clustered with strong support within a distinct subclade closer to the URF root (**Figure 4**, **hashed box**), suggesting that they pre-date 2012. This further suggests that superinfection occurred quite some time before 2012, and also indicates that a majority of intact proviruses predated ART initiation by some time.

**Figure 4.**
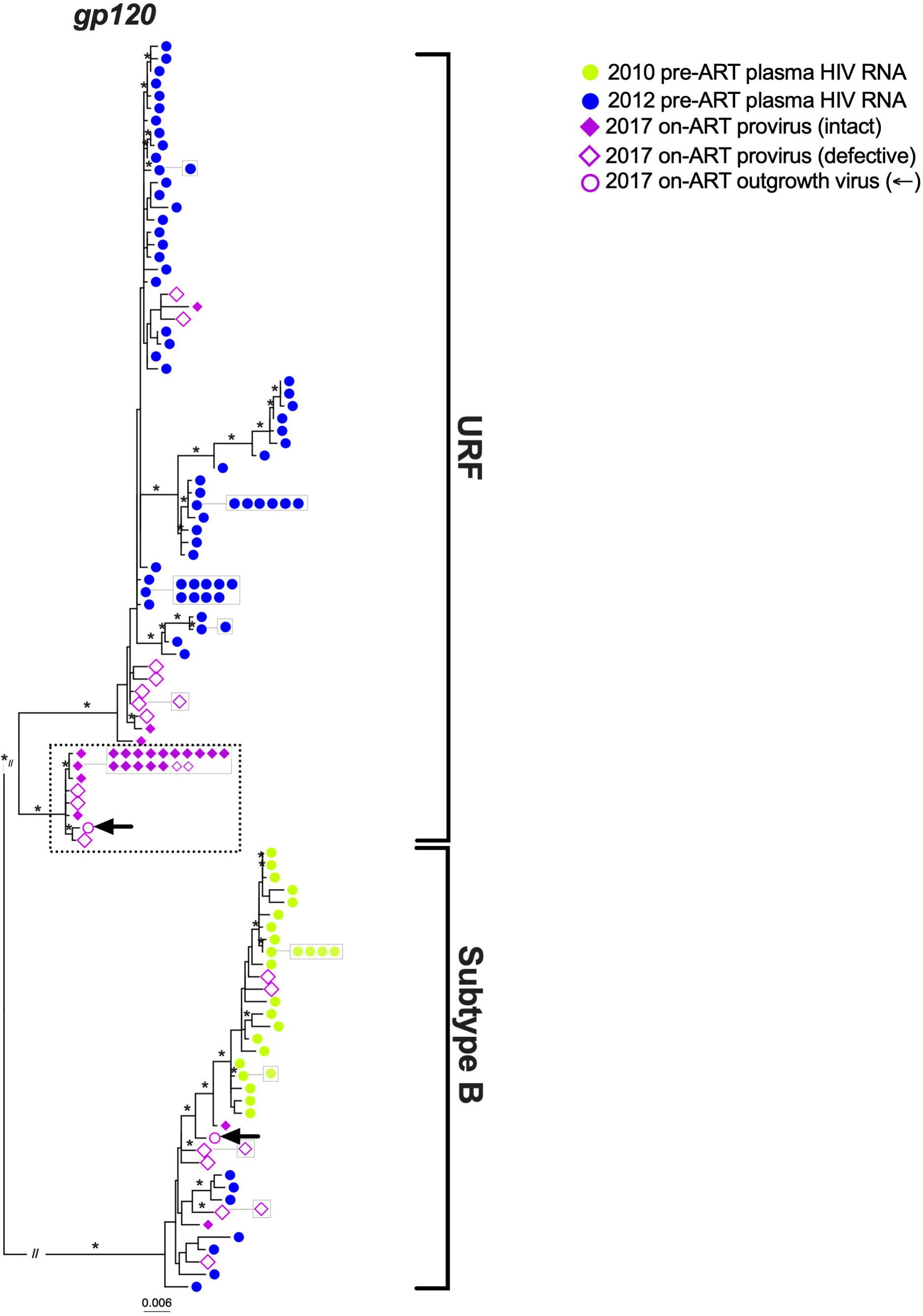
On-ART *gp120* proviral diversity in context of pre-therapy plasma HIV RNA diversity. Maximum-likelihood tree relates pre-ART plasma HIV RNA sequences sampled in 2010 (green circles) and 2012 (blue circles), intact and defective proviruses sampled during ART (solid and open purple diamonds respectively), and outgrowth viruses sampled during ART (open purple circle with arrow). Identical sequences are shown in adjacent boxes. Asterisks identify branches with bootstrap support of ≥90%. Scale is in estimated substitutions per nucleotide site. The tree is midpoint-rooted, with the long subtype-specific branches truncated with a “//” symbol to save space. The hatched box denotes the distinct URF subclade described in the text.

### Initial and superinfecting strains rebounded after ART interruption, with the latter dominating

In 2018 the participant interrupted ART, allowing us to analyze rebound virus evolutionary dynamics. Despite having maintained viral suppression for >6 years, viremia became detectable at 2.8 log_10_ HIV RNA copies/mL after 11 days and rose to 4.8 log_10_ HIV RNA copies/mL by 17 days, which was the first rebound timepoint where plasma was available for analysis (**Figure 5, inset**). Using an overlapping 5-amplicon strategy, we single-genome amplified plasma HIV RNA sequences spanning the entire HIV coding region from samples collected 17, 39 and 53 days post-treatment interruption.

**Figure 5.**
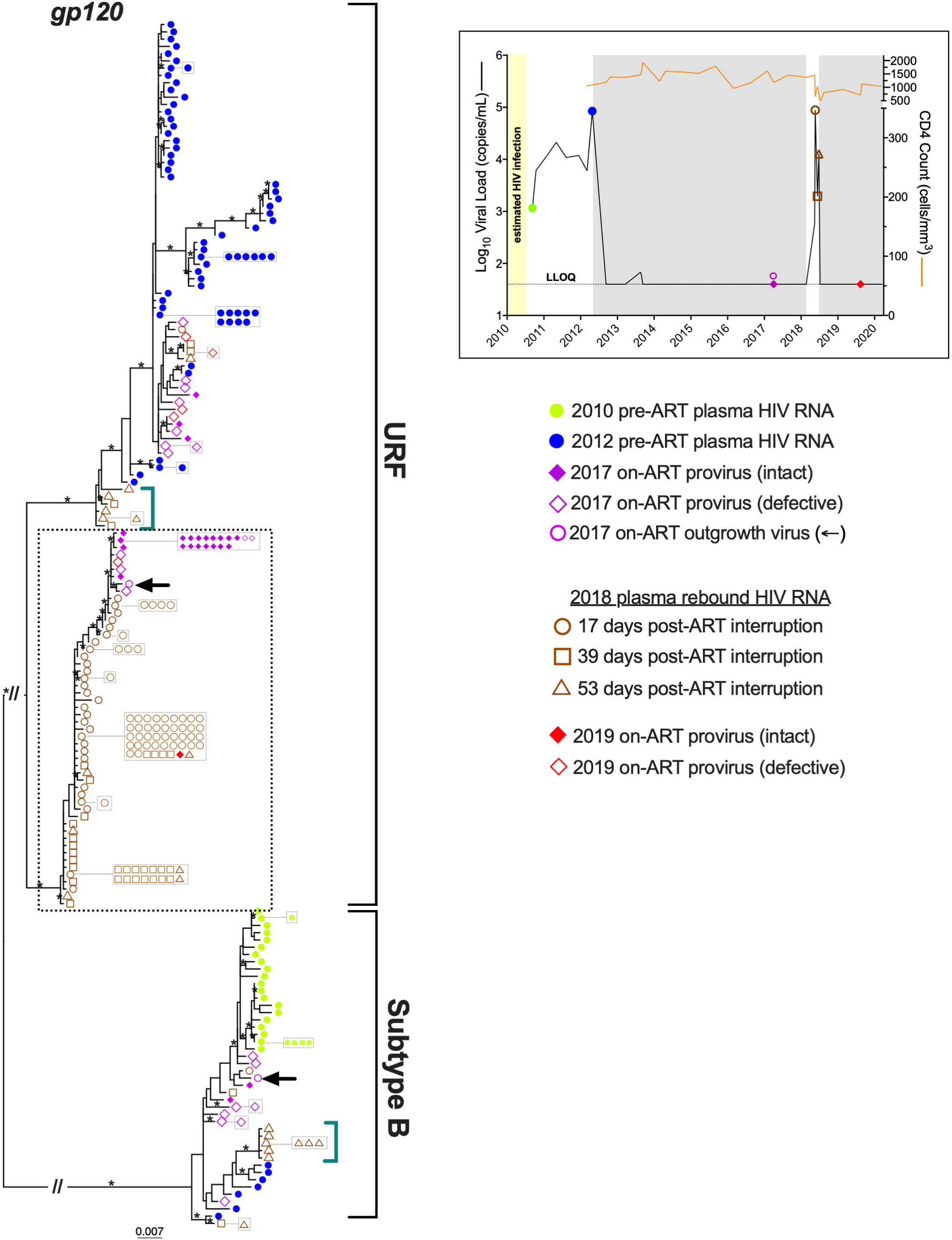
Rebound *gp120* sequences in context of within-host HIV diversity. Maximum-likelihood tree relates pre-ART plasma HIV RNA sequences sampled in 2010 (green circles) and 2012 (blue circles), intact proviruses, defective proviruses and outgrowth viruses sampled in 2017 during ART (solid purple diamonds, open purple diamonds and open purple circles with arrow, respectively), plasma HIV RNA sequences that rebounded during ART interruption (open brown symbols), and intact and defective proviruses sampled in 2019 after ART resumption (solid and open red diamonds). Identical sequences are shown in adjacent boxes. Asterisks identify branches with bootstrap support of ≥90%. Scale is in estimated substitutions per nucleotide site. The tree is midpoint-rooted, with the long subtype-specific branches truncated with a “//” symbol to save space. Hatched box denotes plasma rebound sequences arising from the URF subclade as described in Figure 4. Green brackets denote plasma rebound sequences from later rebound time points that are genetically distinct from those observed at the initial rebound time point. Inset shows the participant’s clinical and sampling history as described in Figure 1.

Despite the underrepresentation of subtype B sequences in the reservoir, genetically diverse HIV *gp120* sequences representing both B and URF strains were present in plasma 17 days after treatment interruption (though URF strains dominated at 99%) (**Figure 5**). This indicates that numerous reservoir cells (or cell clones) reactivated to produce a diverse initial rebounding virus population. Of note, the majority of plasma sequences observed at this point − and throughout the treatment interruption − clustered within the pre-2012 URF subclade that we highlighted in Figure 4, indicating that reservoir cells from this subclade largely fueled the rebound (**Figure 5**, **hashed box**). Descendants of initially rebounding sequences continued to evolve during the treatment interruption, as demonstrated by the isolation of many closely-related *gp120* sequences at the two subsequent timepoints. Numerous identical *gp120* rebound sequences, particularly within the pre-2012 URF subclade, also suggests that rebound was fueled by reactivation of clonally-expanded reservoir cells.

The rebounding sequences outside of this subclade also reveal some insights. Firstly, the only two rebounding sequences isolated at the earliest timepoint that fell outside of this subclade (one URF, one subtype B) clustered relatively closely with intact proviruses isolated from the reservoir the year prior, though branch support values were not strong in all cases (**Figure 5**). This suggests that intact proviruses that were abundant in blood could have fueled the rebound (our ability to isolate these proviruses by single-genome amplification, even once, suggests that they were clonally expanded *in vivo*) (39). Secondly, the subsequent appearance of plasma sequences that were genetically distinct from those sampled earlier in the rebound (see examples in teal brackets in **Figure 5**) are consistent with sequential “waves” of reactivation of different reservoir cells following ART interruption (**Figure 5**). This, and the dominance of the URF during the rebound was confirmed by phylogenetic analysis of *gag*, partial *pol* and *nef* regions (**Figures S1, S4-S5**). No rebound sequence exhibited evidence of recombination between subtype B and URF (**Figures 5, S1, S4-S5** and data not shown).

### Immune escape and drug resistance in the reservoir and during rebound

The participant’s expression of HLA-B*57:03, an allele associated with robust HIV-specific CD8+ cytotoxic T cell responses (23, 24, 40), allowed us to investigate whether rebound sequences were enriched in immune escape mutations compared to the on-ART proviral pool, and whether this differed by subtype. We began with the well-characterized T242N escape mutation within the Gag-TW10 epitope (HXB2 Gag coordinates 240-249) (41, 42). Consistent with escape during untreated infection, T242N was present in 22% of overall subtype B proviruses and 4% of URF proviruses during ART (**Figure 6A**). Notably however, none of the intact subtype B proviruses harbored T242N (all three were susceptible at this residue), though T242N was present as a minority variant among intact URF proviruses (**Figure 6A**). Specifically, the URF virus isolated by viral outgrowth harbored it. At the initial rebound timepoint, 100% of rebounding subtype B sequences harbored the susceptible T242, consistent with their release from T242-containing reservoirs, and no new escape was observed during this period. Also consistent with >95% of URF proviruses harboring the susceptible T242, all initially rebounding URF sequences were T242. But, by 39- and 53-days post-ART interruption, 84% of URF sequences harbored T242N. Notably, all T242N-containing URF rebound sequences fell within a single, limited-diversity subclade where the nearest intact reservoir sequence was the T242N-containing URF outgrowth virus (**Figure S5, hashed box**). This strongly suggests that the outgrowth of T242N-containing URF sequences during the rebound was not due to *de novo* escape, but rather the release of escaped HIV from existing reservoirs that reactivated later. This is also supported by the observation that none of the initial T242 URF rebound variants yielded T242N-containing descendants (in fact, they yielded few descendants at all) (**Figure S5**). Taken together, these observations strongly suggest that TW10-specific memory responses suppressed the initial wave of susceptible T242 variants but not the subsequent wave of T242N-containing ones, ultimately leading to an enrichment in escape variants during the rebound. By contrast, the G248A escape mutation within this same epitope dominated both the proviral pool and the rebound in both subtype B and the URF (≥90%).

**Figure 6.**
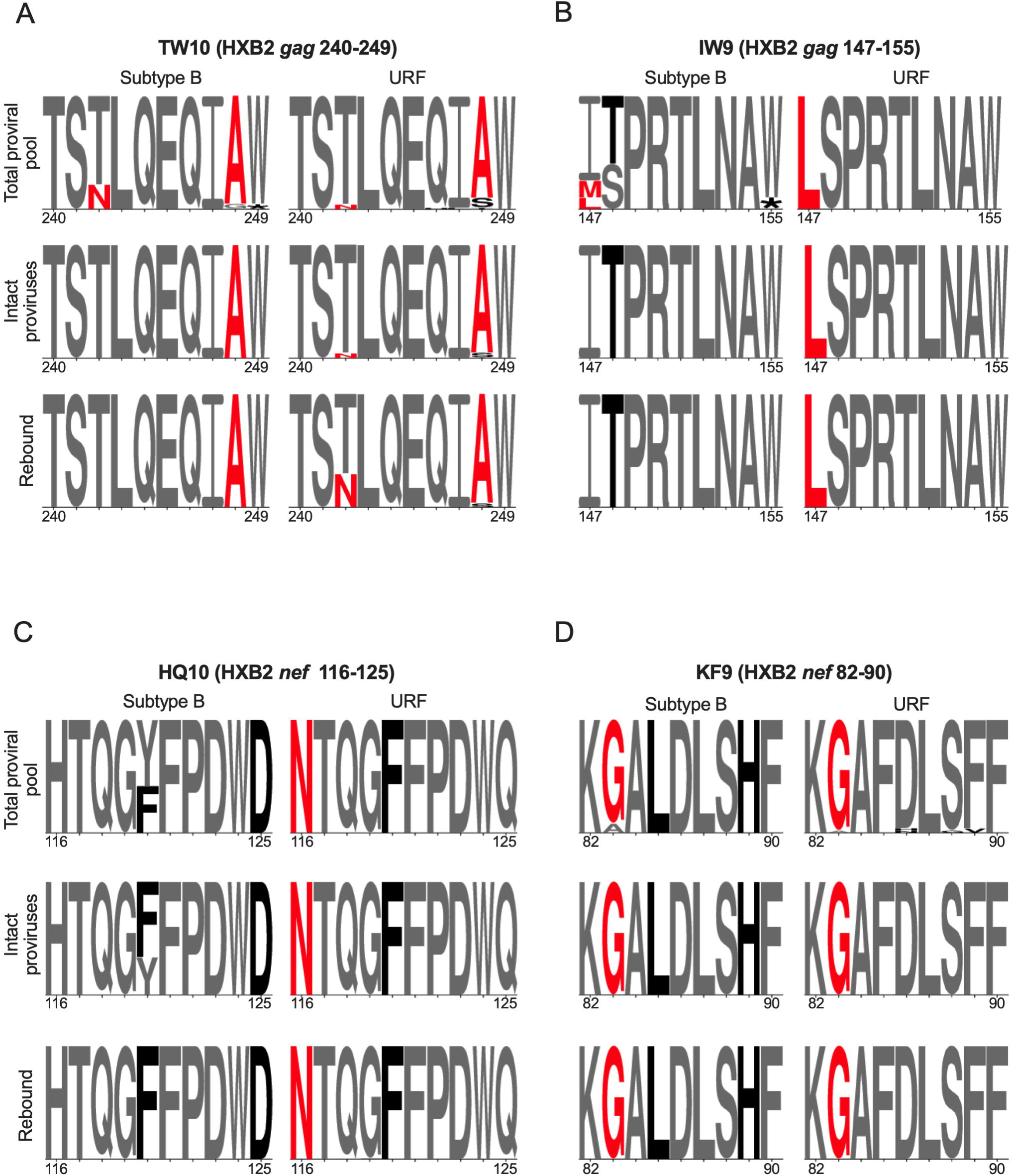
Key HLA-B*57 restricted epitopes in subtype B and the superinfecting URF. Panel A: Subtype B (left) and URF (right) amino acid sequences in the Gag-TW10 epitope, in the total proviral pool excluding hypermutated sequences (top), among intact proviruses (middle), and among rebound viruses (bottom). Letter size denotes amino acid frequency. Red letters denote immune escape mutations. Black letters denote residues that differ from the published epitope sequence. Panels B-D: same as A, but for Gag-IW9, Nef-HQ10, and Nef-KF9 epitopes, respectively.

Similarly, although 21% of overall subtype B proviruses harbored the I147L/M escape mutation within the Gag-IW9 epitope (HXB2 Gag146-155) (43, 44), all three intact proviruses harbored the susceptible I147, which was also observed in 100% of rebounding subtype B sequences, with no subsequent new escape (**Figure 6B**). In contrast, all URF sequences harbored I147L, strongly suggesting that the transmitted/founder URF virus already contained it. Similarly, while all subtype B sequences harbored the susceptible form H116 within the Nef-HQ10 epitope (HXB2 Nef 116-125), 100% of the URF sequences harbored the H116N escape mutation (**Figure 6C**) (45, 46), which is the subtype G consensus at this residue (the URF is subtype G in this region) (47). As such, the URF transmitted/founder virus almost certainly contained this mutation. All intact B and URF proviruses and all rebound viruses harbored the A83G escape mutation (48) within the Nef-KF9 epitope (HXB2 Nef 82-90) (**Figure 6D**), though the existence of a small number of defective proviruses with the susceptible A83 suggests that this mutation was not transmitted, but rather arose through escape during untreated infection. No HLA-B*57 associated escape mutations were found within the Gag-KF11 epitope (HXB2 Gag 162-172) (48), nor in any B*57-restricted Pol epitopes, in either subtype B or the URF.

Clinical drug resistance genotyping performed just prior to ART re-initiation revealed no drug resistance mutations. Of note, this genotype again reported a subtype B classification, though the sequence had a high percentage of amino acid mixtures, consistent with dual infection (not shown). Single-genome sequencing of 36 rebound *pol* sequences confirmed the clinical genotype: no sequences harbored drug resistance mutations except one with a K101E substitution in reverse transcriptase, which confers resistance to certain NNRTIs. K101E was not observed in any prior sequence, and the participant had never received NNRTI-containing ART, so a PCR/sequencing error cannot be ruled out.

### Plasma rebound further enriched URF sequences in the proviral pool

The participant re-initiated ART two days after the final rebound sampling time point, and re-suppressed viremia within two weeks (**Figure 1**). One year later, during which time pVL had been fully suppressed, we isolated 44 near-full length proviruses (39 URF; 5 subtype B) by single-genome sequencing (only 10 million peripheral blood mononuclear cells (PBMC) were available, hence the limited sampling) (**Figure S6)**. Only one provirus, belonging to the URF, was intact. The observation that 89% of 2019 proviruses were the URF, compared to 80% prior to ART interruption, suggests that the ART interruption further skewed reservoir composition towards the URF, which dominated the viral rebound. Also consistent with the rebound event re-seeding the reservoir, T242N frequency among URF proviruses more than doubled to 9% post-interruption, from only 4% pre-interruption. The *gp120* phylogeny also supports reservoir reseeding: 2019 URF proviruses were either identical to, or clustered closely with, plasma rebound sequences (**Figure 5, red diamonds**). This was also apparent in *pol*, *nef* and *gag* (**Figures S1, S4-S5**). Again, all 2019 proviruses were either URF or subtype B, with no evidence of recombination between them (**Figures 5, S1, S4-S5**).

## DISCUSSION

This case confirmed foundational aspects of reservoir dynamics. The recovery of (defective) proviruses related to very early plasma sequences confirms that HIV reservoir seeding begins immediately after transmission (36–38). Nevertheless, the relatively rapid turnover of proviruses − particularly intact ones − during untreated infection means that proviruses persisting during ART are typically enriched in sequences that circulated nearer the time of ART initiation (8, 10–12, 49). This was broadly the case here, though the majority of intact proviruses sampled during ART belonged to a URF subclade that dated to some time before ART initiation, rather than plasma variants that circulated at ART initiation (hatched box, **Figure 4**). The uniqueness of every individual’s reservoir confirms the value of characterizing this heterogeneity to better understand HIV persistence.

This case also shed light on rebound dynamics. Despite the URF dominating the reservoir, both B and URF strains had re-emerged by day 17 following ART interruption, indicating that the first sequences to rebound are not necessarily the most abundant ones in the blood reservoir. The timescale of rebound was very much typical (50), and consistent with reservoir reactivation every 5–8 days (51). The genetic diversity of the initial rebound sequences indicates that multiple diverse reservoir cells (or cell clones) reactivated to release virus (6), while the high frequency of identical *gp120* sequences, particularly within the dominant URF subclade (hashed box in **Figure 5**), suggests that rebound was fueled in part by clonally expanded reservoir cells (52). The continued emergence of rebound sequences that were phylogenetically distinct from the initial rebounding population confirms that diverse reservoirs reactivate continuously during treatment interruption (6). Importantly, the displacement of the initial TW10-susceptible rebounding populations with T242N-containing URF escape variants is consistent with CD8+ cytotoxic T-lymphocyte (CTL) memory responses suppressing the initial wave of susceptible variants but not the subsequent wave of escaped virus emerging from the reservoir (53), underscoring the key role that immune responses play in shaping rebound virus composition and diversity. The outgrowth of T242N-containing URF during the treatment interruption also re-seeded the reservoir, as evidenced by a doubling of T242N frequency among URF proviruses to 9% (from 4%) after the treatment interruption, underscoring a potential risk of such interruptions on the efficacy of future cure strategies that stimulate natural immune responses. Immune escape also provides a plausible explanation for the loss of initial viral control following superinfection: the URF was “pre-adapted” (*i.e.* already harbored B*57:03-associated escape mutations) in key CTL epitopes (54) which would have provided it a substantial fitness advantage over the original subtype B strain.

Recombination is estimated to occur in >70% of superinfection cases (55), so it is interesting that it did not occur in the present case. Different cellular tropism between strains (that could reduce the probability of co-infection events) does not likely explain this, since both B and URF were CCR5-using. Instead, the URF’s higher fitness may have reduced the cellular co-infection opportunities required for recombination (such that any co-infection events would have featured two URF strains) (56), where the incompatibility of the two strains’ dimer initiation signals (DIS) likely further contributed. The DIS is a 6-base palindromic sequence in HIV’s 5’ leader (HXB2 nucleotides 711-716) that mediates the formation of the stable RNA dimers necessary for effective reverse transcription and co-packaging of the genomic RNA (57–59). Mismatches in the DIS restrict HIV intersubtype recombination, and it has been demonstrated that variants containing the canonical subtype B DIS “GCGCGC” (harbored by the infecting subtype B strain) recombine less frequently with those containing the canonical “GTGCAC” of subtypes A, C, F and G (and harbored by the URF) (58, 59). It is also possible that recombinant proviruses arose but were insufficiently fit to compete with the parental strains. Regardless of the mechanism, our observations indicate that specific conditions are required for recombinants to arise and persist.

This study has some limitations. Because pre-ART samples were only available from two time points, we could not determine superinfection timing. Due to very limited plasma quantities, we were unable to study pre-ART immune escape mutations in *gag* and *nef*, and instead focused on pre-ART *pol* and *env* diversity. Similarly, on-ART PBMC quantities were sufficient to genetically characterize proviral diversity but not anti-HIV immune responses. The single plasma rebound sequence with a K101E mutation remains unexplained, and may represent a rare within-host polymorphism, a minority transmitted drug resistance mutation that went undetected despite single-genome amplification, or a RT-PCR/sequencing error.

In conclusion, our findings have implications for HIV clinical management, immune control and cure strategies. This case reminds us that superinfection can go undetected by routine clinical monitoring. While this did not affect the participant’s clinical care, undetected superinfection could theoretically pose risks for data interpretation in HIV cure trials, particularly if the assays used for reservoir monitoring cannot detect diverse HIV subtypes (22). The sequential emergence of genetically diverse HIV sequences following treatment interruption, including immune escape variants that displaced the initial susceptible population, highlights the challenges in predicting rebound virus composition and underscores a key role for immune responses in shaping rebound virus dynamics. Observed evidence that escaped rebound variants subsequently reseeded the reservoir also highlights potential challenges for immune-based HIV cure strategies that harness natural immune responses to eliminate reservoir cells. Finally, our results add to the growing number of molecular studies characterizing HIV reservoirs in non-B subtypes, strengthening the understanding needed to develop an effective HIV cure strategy for all.

## MATERIALS AND METHODS

### Participant Characteristics and Sample Collection

The participant enrolled in a longitudinal HIV cohort study in Toronto, Ontario. The present study analyzes blood samples provided for reservoir characterization in 2017 and 2019, along with archived plasma samples collected during routine clinical care in 2010, 2012 and 2018. The participant provided written informed consent. The cohort study was approved by the institutional ethics boards (IRB) at the University of Toronto, with the IRBs at Providence Health Care/University of British Columbia and Simon Fraser University additionally approving analysis of the participant’s specimens and data.

### Single-Genome HIV amplification and sequencing

HIV RNA *pol* and *gp120* sequences were isolated from plasma as follows. Total nucleic acids were first extracted from 500 μL of plasma on the NucliSENS EasyMag (bioMérieux). For pre-ART plasma, *pol* and *gp120* were single-genome amplified by generating cDNA with HIV-specific primers (**Table S1**), which was then endpoint-diluted such that no more than 30% of subsequent nested PCR reactions would yield amplicons. Amplicons were sequenced on a 3730xl Automated DNA Sequencer (Applied Biosystems) and chromatograms were analyzed using Sequencher (v.5.0, Gene Codes). For rebound plasma, similar approaches were used to single-genome-amplify the entire HIV coding region in five overlapping amplicons spanning *gag-protease*, *protease-reverse transcriptase*, *integrase-vpu*, *gp120*, and *gp41-nef* (primers listed in **Table S1**). These amplicons were sequenced on an Illumina MiSeq. Following bioinformatic removal of primer sequences from MiSeq reads, these were *de novo* assembled using the custom software MiCall (https://github.com/cfe-lab/MiCall), which features an in-house modification of the Iterative Virus Assembler (IVA) (60).

### HIV reservoir quantification

The HIV reservoir was quantified using molecular and viral outgrowth methods. Briefly, the droplet digital PCR (ddPCR)-based intact proviral DNA assay (IPDA) was performed on CD4+ T cells isolated by negative selection from PBMC as previously described (22, 61) where XhoI restriction enzyme (New England Biolabs) was added to each reaction to aid in droplet formation per the manufacturer’s recommendation. Briefly, genomic DNA was isolated from CD4 + T cells using the QIAamp DNA Mini Kit (Qiagen) with precautions to minimize DNA shearing. Intact and total HIV copies/million CD4 + T cells were determined by setting up HIV and human (RPP30 gene) reactions independently in parallel, as described previously (61). In each ddPCR reaction, 7 ng (RPP30) or 700 ng (HIV) of genomic DNA was combined with ddPCR Supermix for Probes (no dUTPs, BioRad), primers (final concentration 900 nM, Integrated DNA Technologies), probe(s) (final concentration 250 nM, ThermoFisher Scientific) and nuclease-free water. Primer and probe sequences are listed in **Table S1**. Droplets were prepared using the Automated Droplet Generator (BioRad). Data were collected on a QX200 Droplet Reader (BioRad) and analyzed using QuantaSoft software (BioRad, version 1.7.4). Replicate wells were merged prior to analysis and intact HIV copies were corrected for DNA shearing as described previously (22, 61).

The quantitative viral outgrowth assay (QVOA) was performed as previously described (22). Briefly, CD4+ T cells were isolated from PBMCs by negative selection and plated in serial dilution at either four or six concentrations (12 replicate wells/concentration, 24-well plates, range 18.6–26.8 million CD4+ T cells total). CD4+ T cells were stimulated with phytohemagglutinin (PHA; 2 µg/mL) and irradiated allogeneic HIV-negative PBMCs were added to further induce viral reactivation. MOLT-4/CCR5 cells were added at 24 hours post-stimulation as targets for viral infection. Culture medium (RPMI 1640 + 10% fetal bovine serum + 1% Pen/Strep + 50 U/mL IL-2 + 10 ng/mL IL-15) was changed every 3 days, and p24 ELISA was run on day 14 to identify virus-positive wells. Culture supernatants from virus-positive wells plated at limiting dilution were immediately used to infect PBMCs from a HIV-negative donor so that the resulting near-full-length provirus could then be amplified and sequenced (see below).

### Near-Full-Length sequencing of HIV proviruses and outgrowth viruses

Proviral DNA was extracted from: 1) CD4+ T cells isolated by negative selection from PBMC collected in 2017, 2) HIV-negative PBMCs inoculated with viral outgrowth sequences isolated at limiting dilution (described above) and 3) directly from PBMC collected in 2019, using the QIAamp DNA Mini Kit (QIAGEN). Near-full length single-genome proviral amplification and sequencing was performed as previously described (9, 62). Briefly, extracted DNA was endpoint-diluted so that no more than ∼25% of the subsequent nested PCR reactions, conducted using Platinum Taq high-fidelity DNA polymerase (Invitrogen), would yield an amplicon (primers in **Table S1**). Amplicons were sequenced on an Illumina MiSeq. For proviruses directly isolated from participant cells, sequencing was performed at the BC Centre for Excellence in HIV/AIDS and reads were *de novo* assembled using MiCall as described above; for those isolated by viral outgrowth, sequencing and genomic assembly was performed at the Massachusetts General Hospital Center for Computational and Integrative Biology core facility. The genomic integrity of each provirus was determined using the open-source software HIV SeqinR, where an “intact” classification requires all HIV reading frames, including accessory proteins, to be intact (63). Sequences with 100% identity across the entire amplicon were considered identical and clonal.

### Sequence alignments, phylogenetic inference and recombination analysis

HIV sequences were codon-aligned using HIVAlign (MAFFT option) (64) hosted on the Los Alamos HIV sequence database (65) or using MUSCLE (66) implemented in Aliview (version 1.28) (67). All alignments were manually inspected using Aliview (version 1.28) (67). Maximum likelihood phylogenies were constructed using IQ-TREE hosted on the IQ-TREE web server (http://iqtree.cibiv.univie.ac.at/) (68) following automated model selection with ModelFinder (69) using a Bayesian information criterion (BIC). Phylogenies were visualized using the R package ggtree (70). Subtyping and estimation of recombination breakpoints for the superinfecting strain was performed using the Recombination Identification Program (RIP) (71) and associated RIP drawing tool (47). Within-host recombination was assessed using RDP4 (25). HIV drug resistance interpretations were performed using the Stanford HIV drug resistance database (version 9.8) (72). Genotypic coreceptor usage predictions were performed using geno2pheno_[coreceptor]_ (73) using the MOTIVATE cutoffs (74).

### Data Availability

GenBank submission is in progress

## ACKNOWLEDGEMENTS

We thank the BC Centre for Excellence in HIV/AIDS clinical laboratory team for support. We gratefully thank the study participant and their HIV care provider, without whom this research would not have been possible. We thank Dr. Marianne Harris, Dr. Mark Hull, Dr. Bruce Ganase, Maria Velasquez and Landon Young for their support with participant recruitment for the cohort shown in Figure 3C.

This work was supported in part by the Canadian Institutes of Health Research (CIHR) through project grant PJT-159625 and focused team grant HB1-164063. This work was also supported in part by CIHR through the Canadian HIV Cure Enterprise (CanCURE) team grant BR4-197730 and the Martin Delaney “REACH” Collaboratory (NIH grant 1-UM1AI164565-01), which is supported by the following NIH co-funding Institutes: NIMH, NIDA, NINDS, NIDDK, NHLBI, and NIAID. FHO was supported by a Ph.D. fellowship from the Sub-Saharan African Network for TB/HIV Research Excellence (SANTHE), a DELTAS Africa Initiative [grant #DEL-15-006]. The DELTAS Africa Initiative is an independent funding scheme of the African Academy of Sciences (AAS)’s Alliance for Accelerating Excellence in Science in Africa (AESA) and supported by the New Partnership for Africa’s Development Planning and Coordinating Agency (NEPAD Agency) with funding from the Wellcome Trust [grant #107752/Z/15/Z] and the UK government. The views expressed in this publication are those of the authors and not necessarily those of AAS, NEPAD Agency, Wellcome Trust or the UK government. NNK was supported by a CIHR Vanier Canada Graduate Scholarship. AS is supported by post-doctoral fellowships from the CIHR and Michael Smith Health Research BC. MCD is supported by a CIHR Doctoral Canada Graduate Scholarship.

F.H.O. and Z.L.B. conceived and designed the study. A.S., N.N.K., W.D., M.C.D., F.Y. and E.B. performed the experiments and collected data. F.H.O., A.S., N.N.K., W.D., V.M. and D.K. analyzed the data. F.H.O. and A.S. visualized the data. M.O., R.M.L., C.K., R.B.J., and C.J.B. provided sample and/or data access. F.H.O. and Z.L.B. wrote the manuscript. All authors critically reviewed the manuscript.

