## Supplemental Figures S1-S6 for "HIV superinfection reveals sequential reservoir reactivation and immune-driven rebound dynamics"

Figure S1

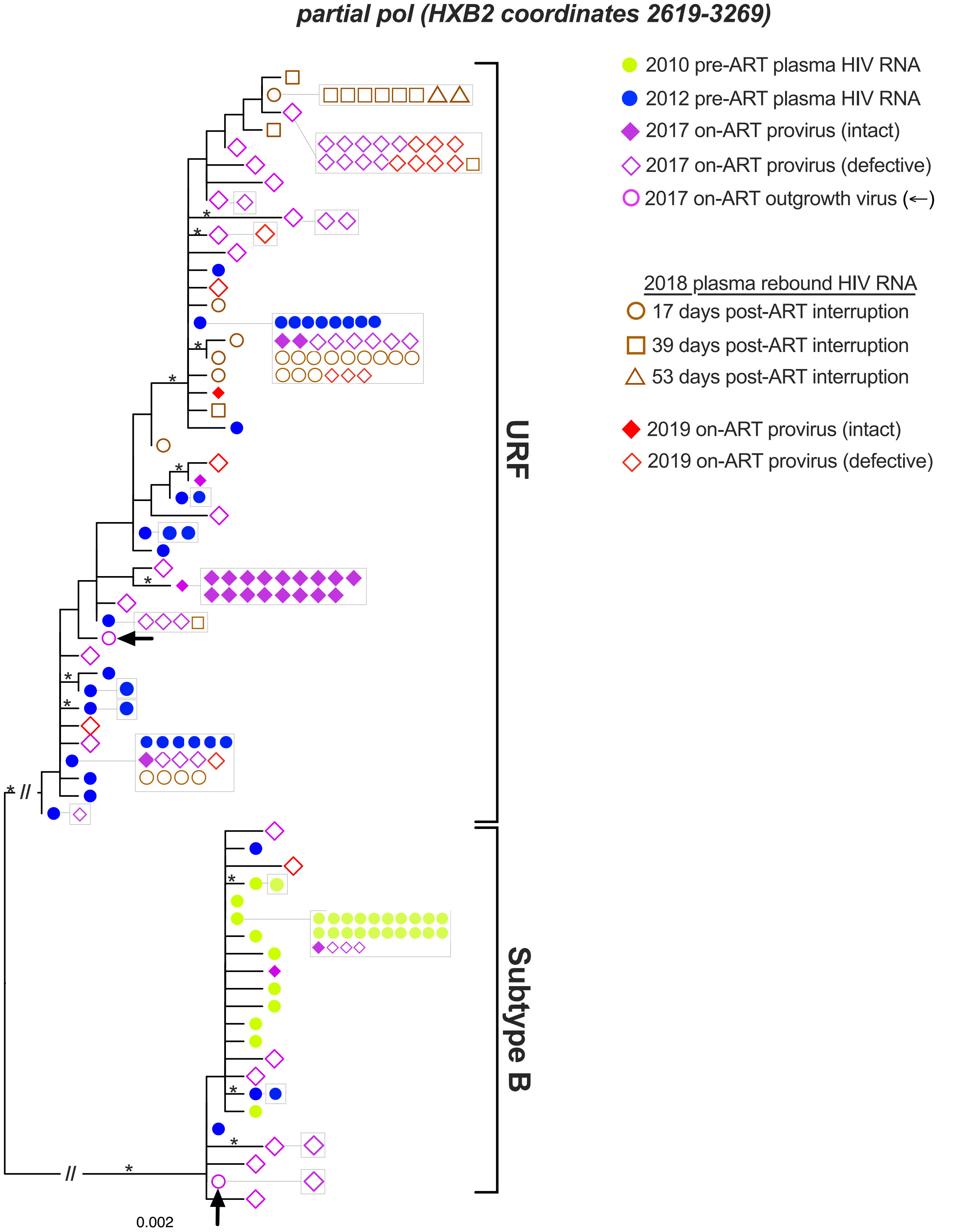

partial pol (HXB2 coordinates 2619-3269)
2010 pre-ART plasma HIV RNA
2012 pre-ART plasma HIV RNA
2017 on-ART provirus (intact)
2017 on-ART provirus (defective)
2017 on-ART outgrowth virus (←)
2018 plasma rebound HIV RNA
17 days post-ART interruption
39 days post-ART interruption
53 days post-ART interruption
2019 on-ART provirus (intact)
2019 on-ART provirus (defective)
URF
Subtype B
0.002

**Figure S1 (previous page). Rebound *pol* sequences in the context of within-host HIV diversity.** Maximum-likelihood tree relates pre-ART plasma HIV RNA sequences sampled in 2010 (green circles) and 2012 (blue circles), intact proviruses, defective proviruses and outgrowth viruses sampled in 2017 during ART (solid purple diamonds, open purple diamonds and open purple circles with arrow, respectively), plasma HIV RNA sequences that rebounded during ART interruption (open brown symbols), and intact and defective proviruses sampled in 2019 after ART resumption (solid and open red diamonds). Identical sequences are shown in adjacent boxes. Asterisks identify branches with bootstrap support of ≥ 90%. Scale is in estimated substitutions per nucleotide site. The tree is midpoint-rooted, with the long subtype-specific branches truncated with a "//" symbol to save space.

Figure S2

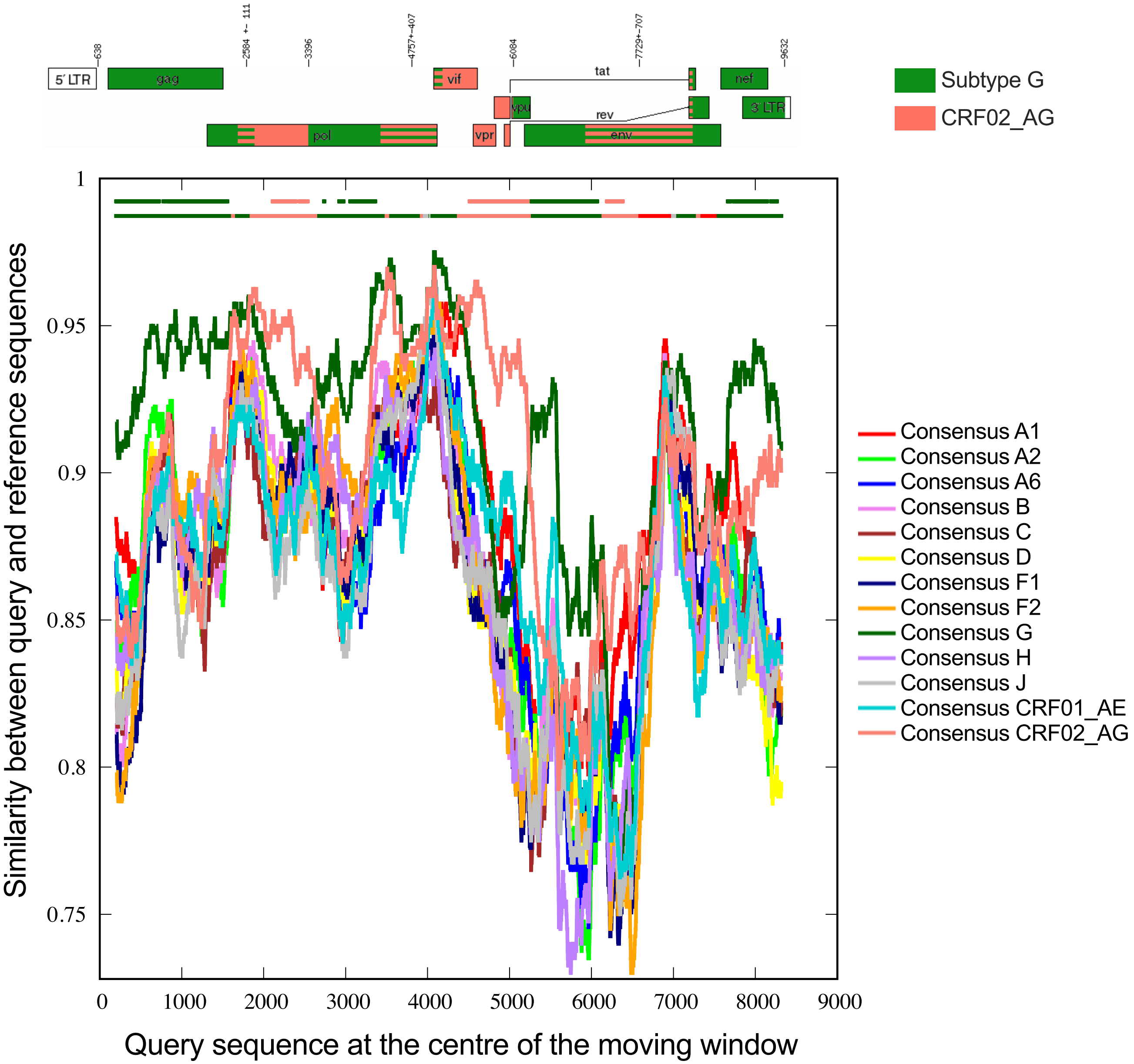

638
2584 +- 111
3396
4757+-407
6084
7729+-707
9632
5' LTR
gag
vif
tat
nef
vpu
rev
3' LTR
pol
vpr
env
Subtype G
CRF02_AG
Similarity between query and reference sequences
Query sequence at the centre of the moving window
1
0.95
0.9
0.85
0.8
0.75
0
1000
2000
3000
4000
5000
6000
7000
8000
9000
Consensus A1
Consensus A2
Consensus A6
Consensus B
Consensus C
Consensus D
Consensus F1
Consensus F2
Consensus G
Consensus H
Consensus J
Consensus CRF01_AE
Consensus CRF02_AG

**Figure S2 (previous page). URF genetic composition.** *Top*: HIV genome map of the superinfecting URF, with subtype G regions shown in green, CRF02_AG regions shown in pink, and hatched sections indicating approximate recombination breakpoints. *Bottom:* Recombinant Identification Program (RIP) plot of the URF genome, using a window size of 400 and a confidence threshold of 95%. Each subtype reference sequence is shown in a unique color. The lines at the top of the plot indicate which subtype reference sequence best matched the superinfecting strain sequence over the genomic region indicated (lower line) and whether this match met the predefined confidence threshold (upper line).

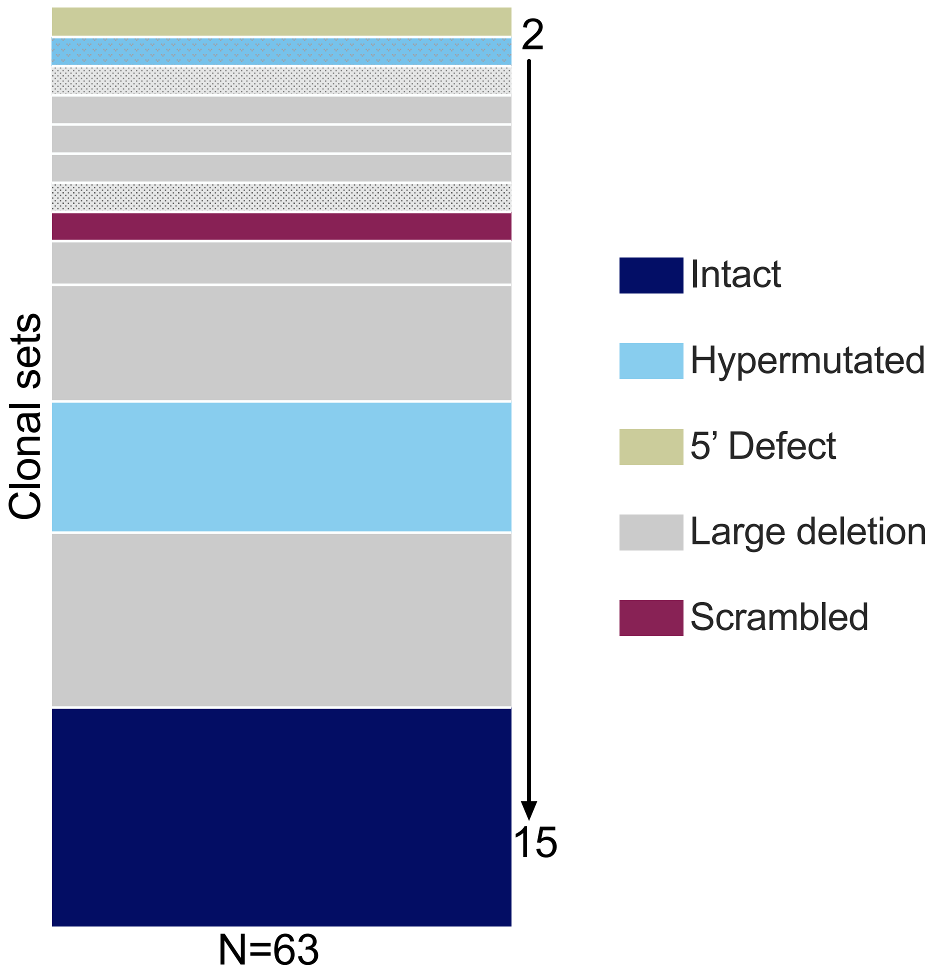
2
15
Clonal sets
N=63
Intact
Hypermutated
5’ Defect
Large deletion
Scrambled

**Figure S3 (previous page). Size and genomic integrity of proviral clones.** All proviruses observed more than once were considered part of a clonal set. We observed a total of 13 clonal sets, which are broken down by clone size (shown by the bar height, where the shortest bar features 2 sequences and the highest bar features 15 sequences) and genomic integrity (shown in different colors). Shading indicates the three clonal sets that are subtype B; all others are the URF.

Figure S4

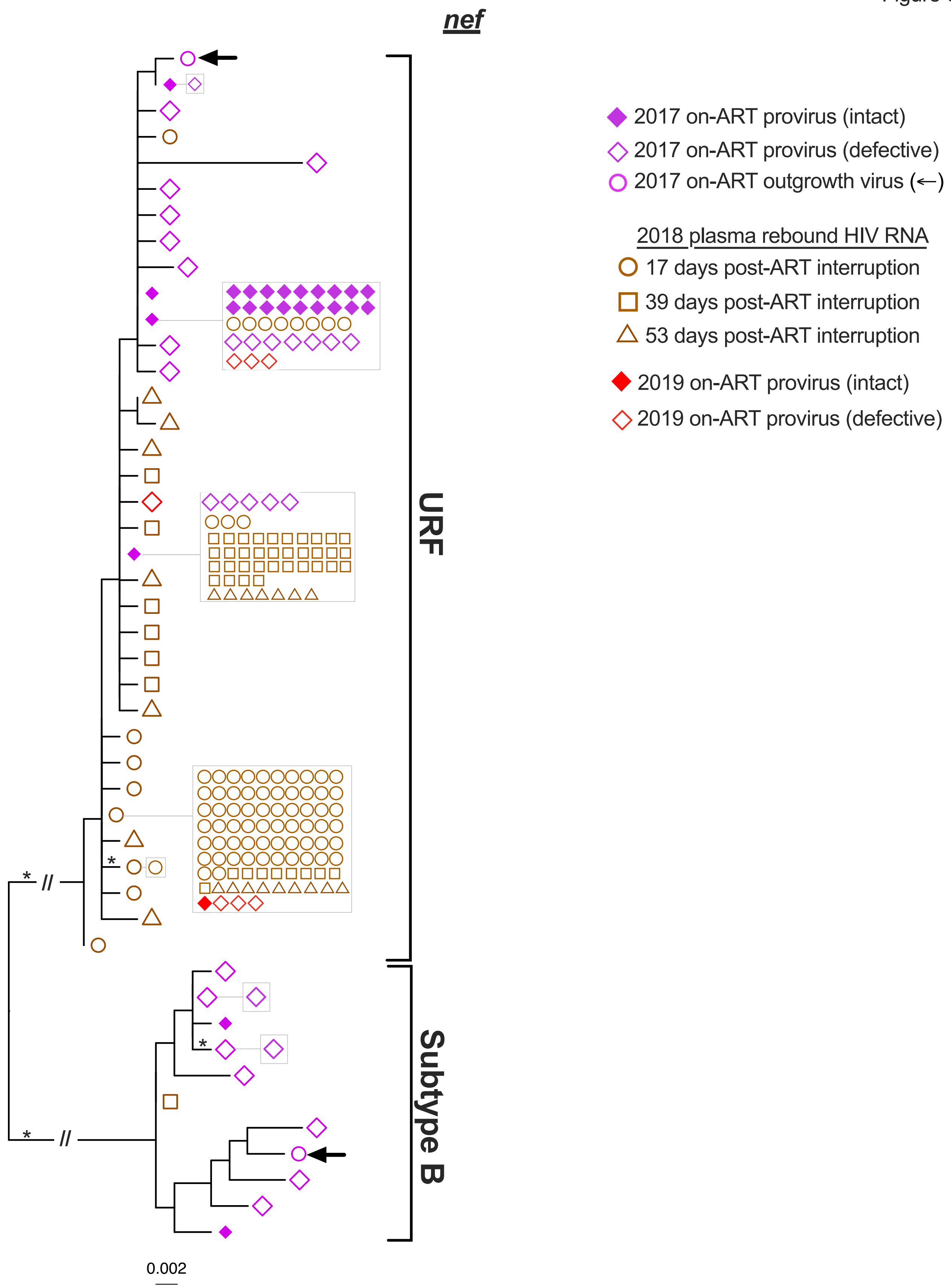
nef
URF
Subtype B
0.002
2017 on-ART provirus (intact)
2017 on-ART provirus (defective)
2017 on-ART outgrowth virus (←)
2018 plasma rebound HIV RNA
17 days post-ART interruption
39 days post-ART interruption
53 days post-ART interruption
2019 on-ART provirus (intact)
2019 on-ART provirus (defective)

**Figure S4 (previous page). Rebound *nef* sequences in the context of on-ART HIV diversity.** Maximum-likelihood tree relates *nef* sequences from intact proviruses, defective proviruses and outgrowth viruses sampled in 2017 during ART (solid purple diamonds, open purple diamonds and open purple circles with arrow, respectively), plasma HIV RNA sequences that rebounded during ART interruption (open brown symbols), and intact and defective proviruses sampled in 2019 after ART resumption (solid and open red diamonds). Identical sequences are shown in adjacent boxes. Asterisks identify branches with bootstrap support of ≥ 90%. Scale is in estimated substitutions per nucleotide site. The tree is midpoint-rooted, with the long subtype-specific branches truncated with a "//" symbol to save space.

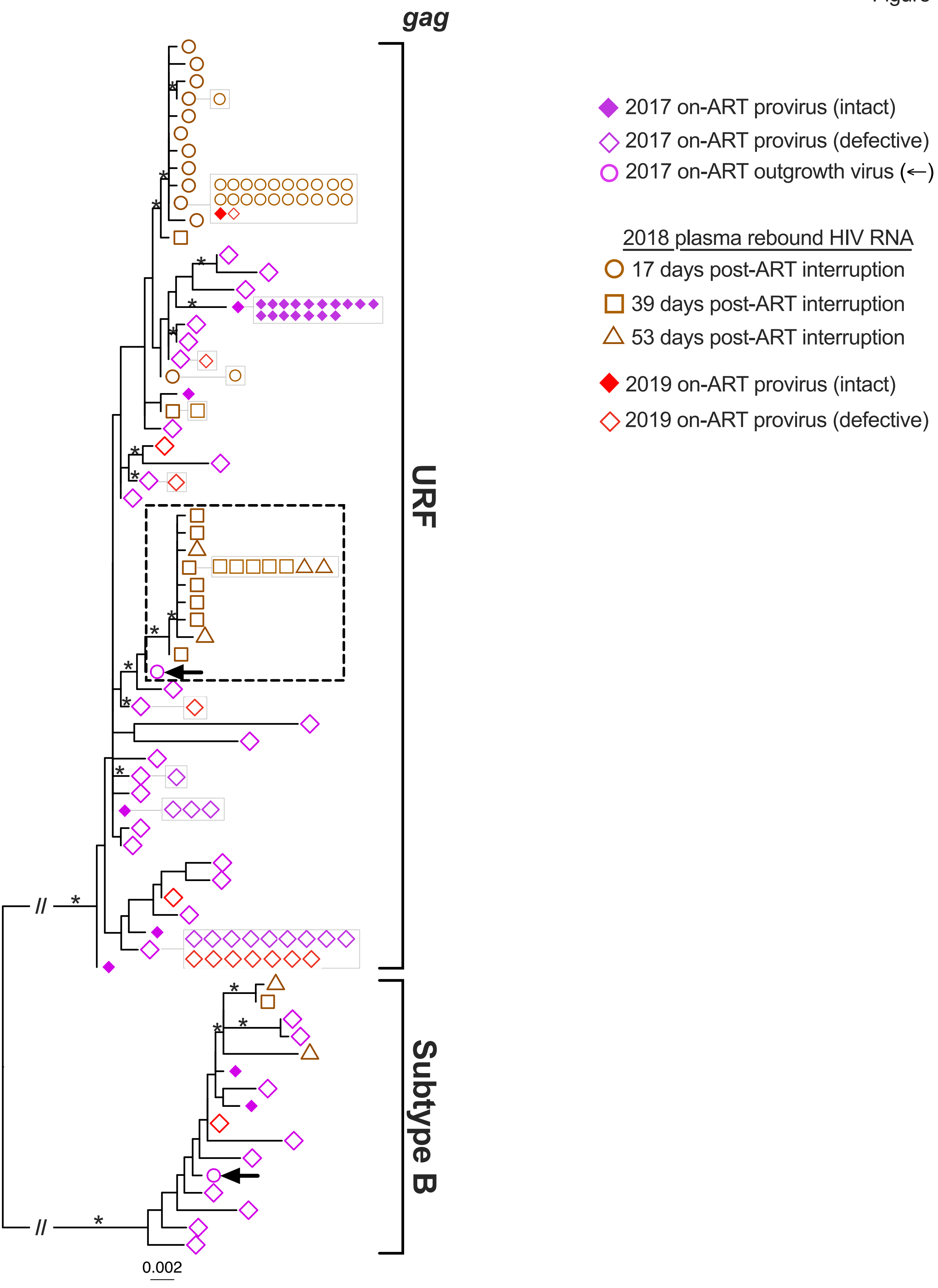
gag
URF
Subtype B
0.002
2017 on-ART provirus (intact)
2017 on-ART provirus (defective)
2017 on-ART outgrowth virus (←)
2018 plasma rebound HIV RNA
17 days post-ART interruption
39 days post-ART interruption
53 days post-ART interruption
2019 on-ART provirus (intact)
2019 on-ART provirus (defective)

**Figure S5 (previous page). Rebound *gag* sequences in the context of on-ART HIV diversity**. Maximum-likelihood tree relates *gag* sequences from intact proviruses, defective proviruses and outgrowth viruses sampled in 2017 during ART (solid purple diamonds, open purple diamonds and open purple circles with arrow, respectively), plasma HIV RNA sequences that rebounded during ART interruption (open brown symbols), and intact and defective proviruses sampled in 2019 after ART resumption (solid and open red diamonds). Identical sequences are shown in adjacent boxes. Asterisks identify branches with bootstrap support of ≥ 90%. Scale is in estimated substitutions per nucleotide site. The tree is midpoint-rooted, with the long subtype-specific branches truncated with a "//" symbol to save space.

Figure S6

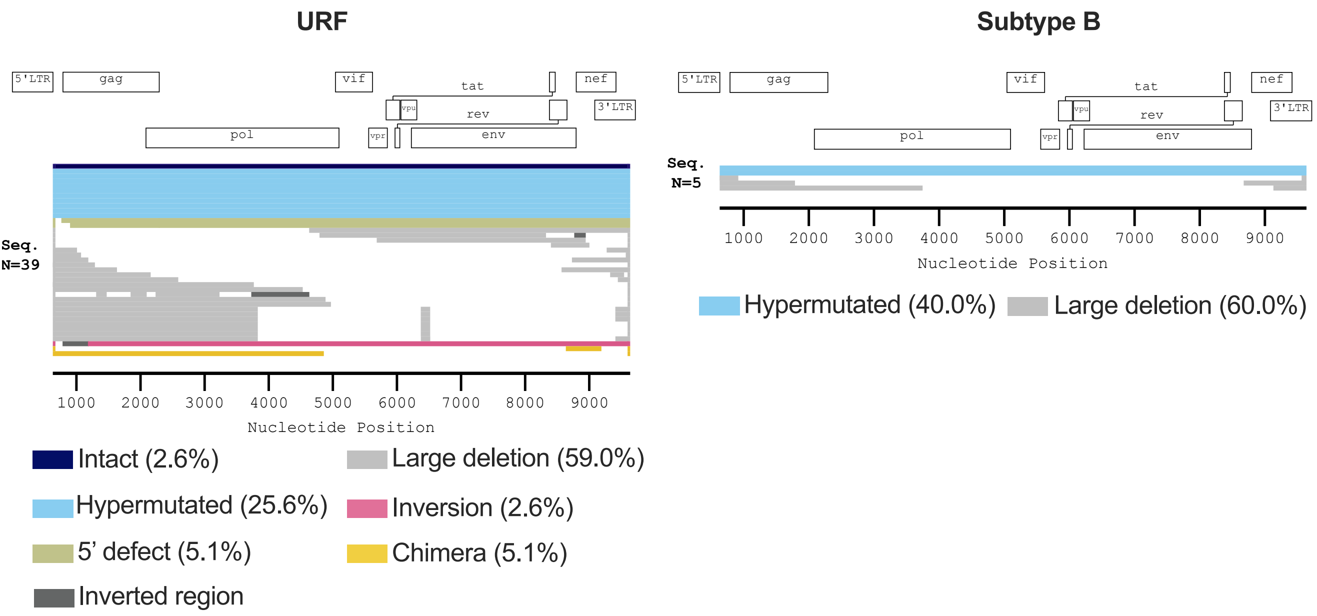
URF
5'LTR
gag
vif
tat
nef
vpu
rev
3'LTR
pol
vpr
env
Seq.
N=39
1000
2000
3000
4000
5000
6000
7000
8000
9000
Nucleotide Position
Intact (2.6%)
Large deletion (59.0%)
Hypermutated (25.6%)
Inversion (2.6%)
5' defect (5.1%)
Chimera (5.1%)
Inverted region
Subtype B
5'LTR
gag
vif
tat
nef
vpu
rev
3'LTR
pol
vpr
env
Seq.
N=5
1000
2000
3000
4000
5000
6000
7000
8000
9000
Nucleotide Position
Hypermutated (40.0%)
Large deletion (60.0%)

**Figure S6 (previous page). Proviral composition one year after ART reinitiation.** Near-full-length URF (left) and B (right) proviral genomes sampled in 2019, one year after ART re-initiation. Proviruses are colored by genomic integrity (white denotes deletions). The frequencies of each sequence type are shown in brackets. The total number of sequences is shown at the left of each plot.
