## Supplemental Table S1 for "HIV superinfection reveals sequential reservoir reactivation and immune-driven rebound dynamics"

**Table S1 - List of primers and probes**

| **Assay** | **PCR reaction** | **Primer name** | **Direction** | **HXB2**  **coordinates** | **Sequence (5' to 3')** |
| --- | --- | --- | --- | --- | --- |
| Near full-length proviral amplification | Outer | 623-Fi(gag) | Forward | 623 - 649 | AAATCTCTAGCAGTGGCGCCCGAACAG |
|  |  | R9662-9686 | Reverse | 9686 - 9662 | TGAGGGATCTCTAGTTACCAGAGTC |
|  | Nested | U5-638F | Forward | 638 - 666 | GCGCCCGAACAGGGACYTGAAARCGAAAG |
|  |  | U5-547R | Reverse | 9632 - 9604 | GCACTCAAGGCAAGCTTTATTGAGGCTTA |
| Near full-length RNA  Amplification | **cDNA** |  |  |  |  |
|  | Primary (full genome) | R9662-9686 | Reverse | 9686 - 9662 | TGAGGGATCTCTAGTTACCAGAGTC |
|  |  | Oligo dT | Reverse | NA | d(TTTTTTTTTTTTTTTTTTTTTTT) |
|  | Alternative (*gag-vpu*) | Env6507R | Reverse | 6529 - 6507 | TCTACCATGTTATTTTTCCACAT |
|  | **Outer** | | | | |
|  | *gag-protease* | 623-Fi(gag) | Forward | 623 - 649 | AAATCTCTAGCAGTGGCGCCCGAACAG |
|  |  | PR.R1 | Reverse | 2782 - 2760 | CTGAAATCTACTAATTTYCTCCA |
|  | *protease-RT* | COMB2 | Forward | 2569 - 2593 | CTGTACCAGTAAAATTAAAGCCAGG |
|  |  | IN4801R | Reverse | 4801 - 4776 | ATCCCCCCTTTTCTTTTAAAATTGTG |
|  | *int-vpu* | RT4035F | Forward | 4035 - 4060 | GTAACAGACTCACAGTATGCATTAGG |
|  |  | Env6507R | Reverse | 6529 - 6507 | TCTACCATGTTATTTTTCCACAT |
|  | *gp120* | SC02F.1 | Forward | 5956 - 5980 | CTTAGGCATCTCCTATGGCAGGAAG |
|  |  | C0602 | Reverse | 7817 - 7786 | GCCCATAGTGCTTCCTGCTGCTCCCAAGAACC |
|  | *gp41-nef* | GP41Fo | Forward | 7626 - 7648 | TTCAGACCTGGAGGAGGAGATAT |
|  |  | R9662-9686 | Reverse | 9686 - 9662 | TGAGGGATCTCTAGTTACCAGAGTC |
|  | **Nested** | | | | |
|  | *gag-protease* | GS1F | Forward | 688 - 709 | GACGCAGGACTCGGCTTGCTGA |
|  |  | ProC- | Reverse | 2724 - 2697 | GAGTATTGTATGGATTTTCAGGCCCAAT |
|  | *protease-RT* | CP2F | Forward | 2610 - 2635 | GTTAAACAATGGCCATTGACAGAAGA |
|  |  | IN4678R | Reverse | 4678 - 4654 | ACTCCTTGACTTTGGGGATTGTAGG |
|  | *int-vpu* | INT4195F | Forward | 4173 - 4195 | ATTGGAGGAAATGAACAAGTAGA |
|  |  | ENV6437R | Reverse | 6459 - 6437 | GGTCTGTGGGTACACAGGCATGT |
|  | *gp120* | ENVF.1 | Forward | 6203 – 6224 | GAAAGAGCAGAAGACAGTGGCA |
|  |  | CD4R | Reverse | 7675 - 7652 | TATAATTCACTTCTCCAATTGTCC |
|  | *gp41-nef* | GP41Fi | Forward | 7652 - 7674 | GGACAATTGGAGAAGTGAATTAT |
|  |  | R9604-9632 | Reverse | 9632 - 9604 | GCACTCAAGGCAAGCTTTATTGAGGCTTA |
| Drug Resistance Genotyping; Protease/RT | cDNA | RT3.1 | Reverse | 3859-3830 | GCTCCTACTATGGGTTCTTTCTCTAACTGG |
|  | Outer | 5CP1 | Forward | 1981-2008 | GAAGGGCACACAGCCAGAAATTGCAGGG |
|  |  | RT3.1 | Reverse | 3859-3830 | GCTCCTACTATGGGTTCTTTCTCTAACTGG |
|  | Nested | 2.5 | Forward | 2011-2039 | CCTAGGAAAAAGGGCTGTTGGAAATGTGG |
|  |  | RT3798R | Reverse | 3798-3777 | CAAACTCCCACTCAGGAATCCA |
| Intact Proviral DNA Assay | RPP30 (human) | RPP30Fwd | Forward | NA | GATTTGGACCTGCGAGCG |
|  | RPP30 (human) | RPP30Probe | Probe | NA | VIC^a^-CTGACCTGAAGGCTCT-MGBNFQ^b^ |
|  | RPP30 (human) | RPP30Rev | Reverse | NA | GCGGCTGTCTCCACAAGT |
|  | RPP30-Shear (human) | ShearFwd | Forward | NA | CCATTTGCTGCTCCTTGGG |
|  | RPP30-Shear (human) | ShearProbe | Probe | NA | 6-FAM^c^-AAGGAGCAAGGTTCTATTGTAG-MGBNFQ |
|  | RPP30-Shear (human) | ShearRev | Reverse | NA | CATGCAAAGGAGGAAGCCG |
|  | HIV *psi* | GagFwd | Forward | 692 - 711 | CAGGACTCGGCTTGCTGAAG |
|  | HIV *psi* | GagProbe | Probe | 758 - 740 | 6-FAM-TTTTGGCGTACTCACCAGT-MGBNFQ |
|  | HIV *psi* | GagRev | Reverse | 797 - 775 | GCACCCATCTCTCTCCTTCTAGC |
|  | HIV *env* | EnvFwd | Forward | 7736 - 7759 | AGTGGTGCAGAGAGAAAAAAGAGC |
|  | HIV *env* | EnvProbe | Probe | 7781 - 7796 | VIC-CCTTGGGTTCTTGGGA-MGBNFQ |
|  | HIV *env* | EnvRev | Reverse | 7851 - 7832 | GTCTGGCCTGTACCGTCAGC |
|  | HIV *env* | HypermutProbe | Probe | 7781 - 7798 | CCTTAGGTTCTTAGGAGC-MGBNFQ |

^a^VIC = 2′-chloro-7′phenyl-1,4-dichloro-6-carboxy-fluorescein

^b^MGBNFQ = *3’* Minor groove binder; NFQ = Nonfluorescent quencher

^c^6-FAM = carboxyfluorescein
